# Recent evolution of extreme cestode growth suppression by a vertebrate host

**DOI:** 10.1101/091942

**Authors:** Jesse N. Weber, Natalie C. Steinel, Kum Chuan Shim, Daniel I. Bolnick

## Abstract

Parasites can be a major cause of natural selection on hosts, which consequently evolve a variety of strategies to avoid, eliminate, or tolerate infection. When ecologically similar host populations present disparate infection loads, this natural variation can reveal immunological strategies underlying adaptation to infection and population divergence. For instance, the tapeworm *Schistocephalus solidus* persistently infects between 0% to 80% of threespine stickleback (*Gasterosteus aculeatus*) in lakes on Vancouver Island. To test whether these heterogeneous infection rates are due to evolved differences in immunity, we experimentally exposed lab-reared fish from high-and low-infection populations, which are not known to differ in natural exposure risk, to controlled doses of *Schistocephalus.* We observed heritable between-population differences in several immune traits: fish from the naturally uninfected population initiated a stronger granulocyte response to *Schistocephalus* infection, and their granulocytes constitutively generated threefold more reactive oxygen species (ROS). Despite these immunological differences, *Schistocephalus* was equally successful at establishing initial infections in both host populations. However, the low-infection fish dramatically suppressed tapeworm growth relative to high-infection fish, and parasite size was intermediate in F1 hybrid hosts. Our results show that stickleback recently evolved heritable variation in their capacity to suppress helminth growth. Comparative data from many from natural populations indicate that growth suppression is widespread but not universal and, when present, is associated with reduced infection prevalence. Host suppression of helminth somatic growth may be an important immune strategy that aids in parasite clearance, or in mitigating the fitness costs of persistent infection.

**Significance:** Large parasites remain a persistent source of morbidity and mortality in humans, domesticated animals, and wildlife. Hosts are subject to strong natural selection to eliminate or tolerate these parasite infections. Here, we document the recent evolution of a striking form of resistance by a vertebrate host (threespine stickleback) against its cestode parasite (*Schistocephalus solidus).* After Pleistocene glacial retreat, marine stickleback colonized freshwater lakes, encountered *Schistocephalus*, and evolved varying levels of resistance to it. We show that a heavily-and a rarely-infected population of stickleback have similar resistance to *Schistocephalus* colonization, but rarely-infected fish suppress parasite growth by orders of magnitude. These populations represent ends of a natural continuum of cestode growth suppression, which is associated with reduced infection prevalence.

## Introduction

Helminth parasites (trematodes, nematodes, cestodes) currently infect approximately 24% of humanity (1), undermine agricultural productivity (2, 3), and threaten conservation of some wild populations (4). These social, economic, and environmental costs motivate substantial interest in how vertebrate hosts combat helminth infection. Helminths often severely reduce their hosts’ fitness, and therefore represent a strong source of natural selection on their hosts (5–7). In response, vertebrates have evolved a complex repertoire of innate and adaptive immune responses that serve to detect and eliminate infections, or to promote tolerance of successful infections.

Despite vertebrates’ sophisticated immune systems, helminths remain common and persistent, often establishing infections that last months or years. Parasites’ continued success may be attributed to (i) hosts’ failure to evolve effective resistance due to insufficient time or genetic diversity (8), (ii) trade-offs that constrain hosts’ ability to evolve costly immune traits (9–12), or (iii) parasites’ counter-adaptations to undermine host immunity (13). These factors may apply unequally across host populations, due to differences in host-parasite encounter rates, host genetic diversity, co-evolutionary time, ecological costs of immunity, or parasite genotypes (14). As a result, some hosts may be substantially more resistant or tolerant than others.

Natural variation in host immunity provides an opportunity to elucidate vertebrates’ evolving strategies to limit helminth infections. This can be done by surveying diverse natural populations to identify instances of exceptionally high or low parasite prevalence and testing whether these differently-infected populations vary with respect to immune phenotypes. Finally, we can evaluate whether genetic differences in resistance are responsible for the natural variation in infection rates. This approach is exciting because it can identify immune strategies that natural selection has favored to mitigate parasite infections. Unlike traditional immunogenetics, which uses mutagenesis to create phenotypic aberrations [frequently involving loss-of-function mutations (15)], or forward genetic approaches to map causative loci (16), natural selection scans entire populations for many generations to find beneficial resistance traits.

Here, we document wide-ranging variation in parasite abundance (from 0% to 80% prevalence) in wild populations, and show that this variation is associated with heritable differences in immune response and the evolutionary gain of an underappreciated form of resistance: parasite size suppression. Threespine stickleback *(Gasterosteus aculeatus)* inhabit brackish and freshwater habitats throughout coastal north temperate regions. After Pleistocene deglaciation (~12,000 years ago), marine stickleback colonized many replicate freshwater lakes on Vancouver Island and elsewhere. This colonization brought marine fish into contact with a freshwater parasite, the cestode *Schistocephalus solidus.* Because *S.solidus* eggs do not hatch in brackish water (17), marine stickleback are rarely infected by this cestode, and therefore have not evolved effective resistance (18). Newly established freshwater populations encountered *S.solidus* at a higher rate, resulting in reduced survival and fecundity (19, 20), and selection for increased resistance. Independently-colonized lake populations evolved parallel gain of resistance (18). Here, we show that this parallel evolution is incomplete: lake populations differ with respect to immune responses, and parasite resistance, because some populations evolved effective suppression of cestode growth.

## Results

### Tapeworm Infection Prevalence Differs Among Natural Stickleback Populations

Samples of stickleback from 50 lakes on Vancouver Island (Table S1) revealed among-population variation in *S. solidus* infection prevalence, spanning at least two orders of magnitude and persisting for a decade (Fig 1, *χ*^2^=4629.3, df=49, P<0.0001). To experimentally test for variation in stickleback resistance to *S.solidus*, we focused on two lakes that bracket the range of infection prevalence. Gosling Lake stickleback (GOS) had the second highest infection prevalence in our sample (50-80%, depending on the year). In contrast, no infections were observed in Roberts Lake stickleback (ROB; Fig 1; N=1480 fish).

**Figure 1:**
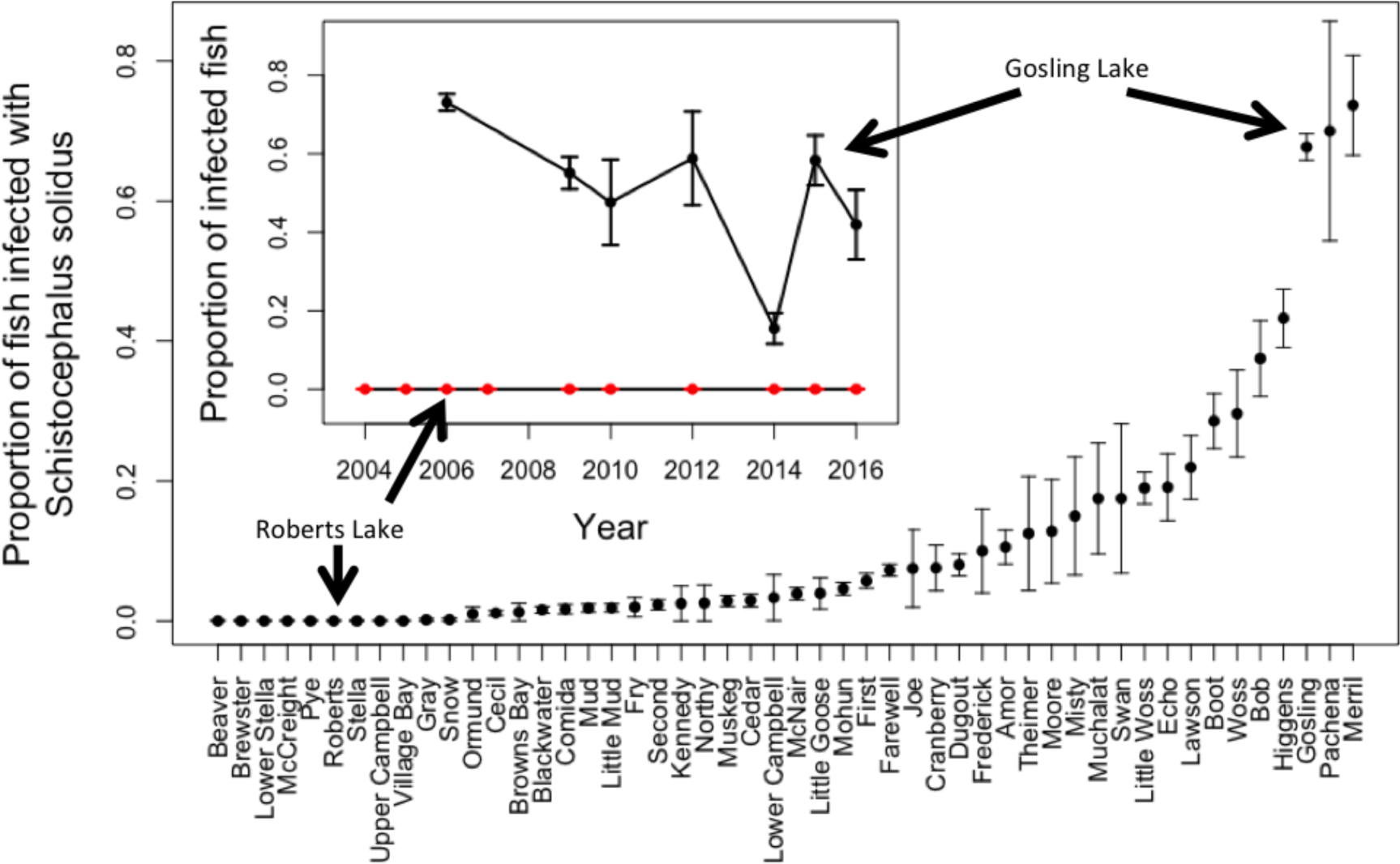
Natural variation in *S.solidus* prevalence among 50 lakes on Vancouver Island, British Columbia. Sample year, sample size, and location details are in Table S1. Error bars are SEs of each sample. The inset presents data from two focal populations, Roberts and Gosling Lakes, over a roughly 10-year time span.

Ecological differences between the two lakes are minimal. Gosling and Roberts Lakes are only 17 km apart, both moderately large (60 and 160 hectares, respectively) and similarly deep (40 and 53m max depth). Both lakes contain large pelagic zones in which stickleback forage on copepods *(S.solidus'* first host) at similar rates [averaging 0.444 and 0.457 cyclopoid copepods per fish per day, respectively (21); Poisson GLM P=0.9707 for 27 and 35 fish]. Both lakes host at least one breeding pair of loons (the terminal host) per summer, which we have observed foraging in nearby lakes where *S.solidus* is common (DIB observation).

Consequently, there is no currently-known difference in *S.solidus* exposure risk between stickleback in Gosling and Roberts Lakes.

### Divergence in Immune Responses

We used a common garden breeding experiment to test whether GOS and ROB stickleback evolved divergent immune phenotypes. We crossed wild-caught adults from GOS and ROB to generate 51 families, which included both pure-population fish and F1 hybrids with GOS or ROB dams (GxR or RxG, respectively). The eggs were hatched and reared to adulthood in laboratory aquaria. We also bred *S.solidus* plerocercoids obtained from infected fish from three lakes in British Columbia. Approximately five fish per family were fed copepods containing infective *S. solidus* procercoids (Fig S1). One to two siblings per family were fed uninfected copepods as a control (Fig S1; data are deposited at dryad.org). Approximately forty-two days post-exposure, we measured host immune phenotypes to test for constitutive and infection-induced differences between stickleback populations. Here, we focus on two traits previously associated with defense against macroparasites: the ratio of granulocytes to lymphocytes in head kidney (HK) primary cell cultures, and the magnitude of reactive oxygen species (ROS) produced by granulocytes.

Granulocytes constituted a lower proportion of HK cells in ROB fish (5.5% less than GOS fish; t=2.31, P=0.03). Despite this lower initial frequency, ROB fish responded to infection by increasing granulocyte proportions (16.3% higher than uninfected ROB fish; t=5.40, P≤0.001; Fig 2A). Infection had no effect on GOS fish, resulting in an interaction between fish population and infection status (Table S2). A similar interaction occurred in F1 hybrids: infection did not affect granulocyte frequency in GxR hybrids, but their frequency increased 12.8% in infected RxG fish, relative to uninfected siblings (t=3.37, P=0.0009; Fig 2A). This asymmetry in response to infection by reciprocal F1 hybrids strongly suggests that a maternal effect in ROB stickleback drives the observed phenotype.

**Figure 2.**
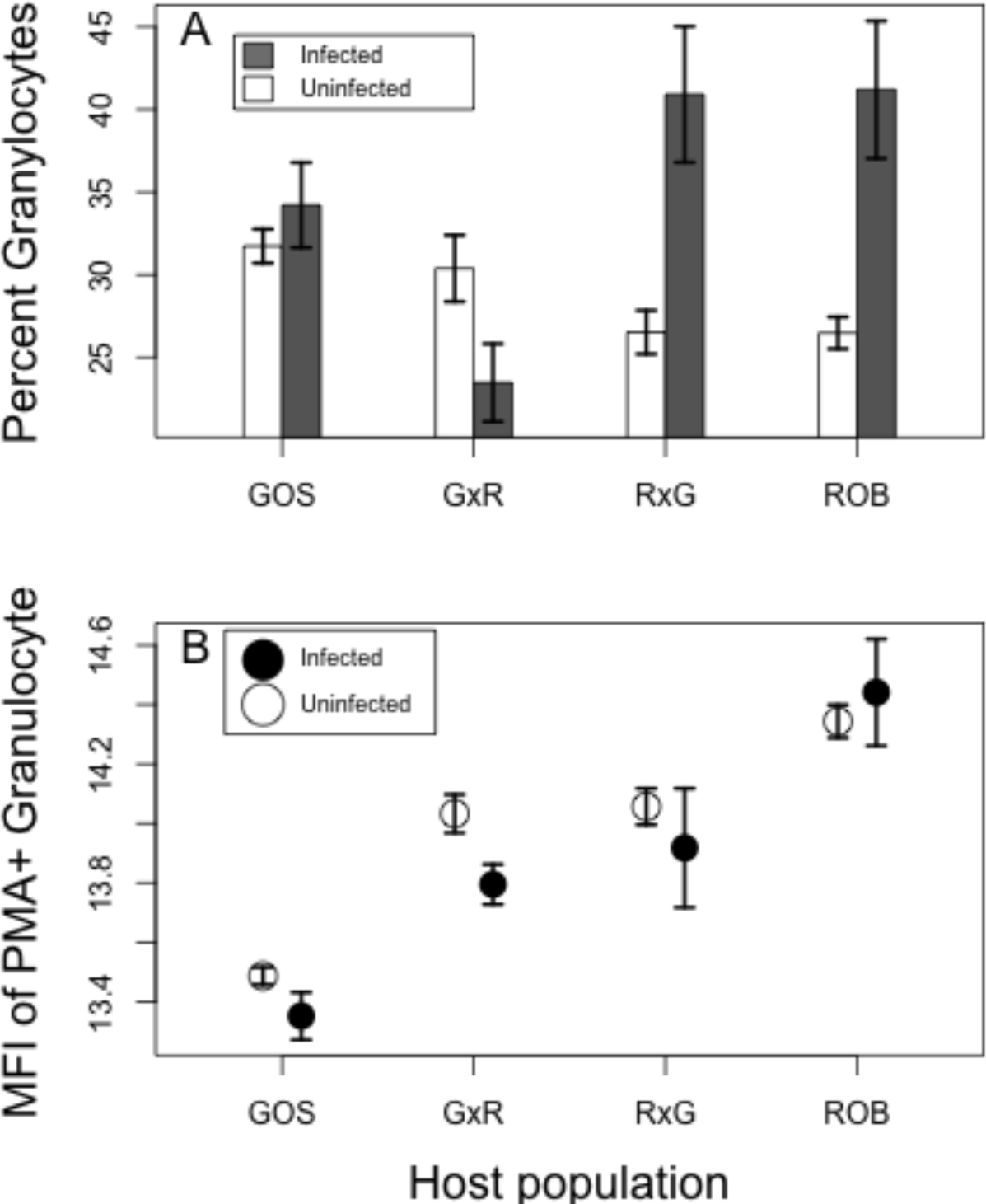
Variation in stickleback immunity, as measured by (A) the percentage of granulocytes in HK primary cell culture, and (B) the median fluorescence intensity (MFI) of PMA-stimulated granulocytes. Error bars are one standard error intervals above and below the mean. Sample size of uninfected and infected fish (respectively) in each population: GOS=95 and 16; GxR=43 and 10; RxG=57 and 7; ROB=105 and 9.

The relative abundance of granulocytes also differed between the sexes (4.7% less in males; t=4.03, P=8.4e-5; Fig. S2), and varied among stickleback families within the two populations (random family effect, Table S2). Thus, there is segregating genetic variation for this trait within both populations. Granulocyte relative abundance was independent of cestode genotype (Table S2), and unaffected by experimental exposure to *S.solidus* (Fig. S3).

The production of reactive oxygen species (ROS) was quantified by measuring the median fluorescence intensity (MFI) of HK cells stained with Dihydrorhodamine-123 (DHR). DHR is a cell-permeable fluorophore that fluoresces in the presence of peroxide and peroxynitrite (two forms of ROS). For each fish we measured: (1) baseline ROS production in unstimulated granulocytes and (2) induced ROS production in Phorbol 12-myristate 13-acetate (PMA)-stimulated granulocytes. Here, we focus on the second metric (see Fig. S4 for baseline ROS). PMA-stimulated ROB cells generated 2.7-fold greater MFI than GOS cells (main effect of population on ln-transformed MFI, t=7.77, P<7.8e-9; Fig. 2B). ROS production by F1 hybrids’ granulocytes was intermediate between ROB and GOS stickleback (Fig. 2B). Both hybrid crosses produced more ROS than GOS fish (both P≤0.0034 in post-hoc pairwise Tukey tests). GxR and RxG families generated equivalent ROS to each other (P=0.988) and less than ROB fish (P=0.034 and 0.061, respectively). Because this trait did not differ between infected and uninfected (control or experimental) stickleback, nor was there an interaction between fish population and infection status (Table S2), we infer that ROS production differs constitutively between GOS versus ROB fish. Similar to granulocyte frequency, there is segregating genetic variation in ROS production within populations (random family effect, Table S2). Neither fish sex nor cestode genotype had detectable effects on granulocyte ROS production. Overall, we conclude that GOS and ROB stickleback display heritable differences in immune responses. Some differences are constitutive (i.e., ROS levels), others are induced by infection (i.e., % granulocytes).

### Recently-Evolved Differences in Resistance

GOS and ROB stickleback did not differ significantly in the frequency or intensity of laboratory infections (Fig 3A&B). Infection rates in GOS and ROB fish, respectively, were 20.0% and 11.4% (with standard errors of 4.5% and 3.6%; binomial GLM, Z=-1.47, P=0.14). Including random effects of host family did not improve model fit (AIC=142.3 and 140.4 with and without family), suggesting little effect of segregating genetic variation within populations. Although parasite genotype did not significantly affect infection success rate (P=0.81), there was a marginally significant interaction between host and parasite genotype (P=0.09; Fig. S5). Infected GOS and ROB fish carried equivalent numbers of cestodes (1.75 versus 1.78 out of 5 cestodes provided per fish, binomial GLM, Z=0.06, P=0.95). We conclude that the absence of *S.solidus* from wild Roberts Lake fish is not attributable to any greater resistance to colonization by *S.solidus.* This conclusion implies that the separately-colonized lake populations independently evolved quantitatively similar resistance to *S.solidus* establishment, relative to their highly susceptible marine ancestors (18). This represents a rarely documented instance of convergently-evolved helminth immunity in a wild vertebrate.

**Figure 3.**
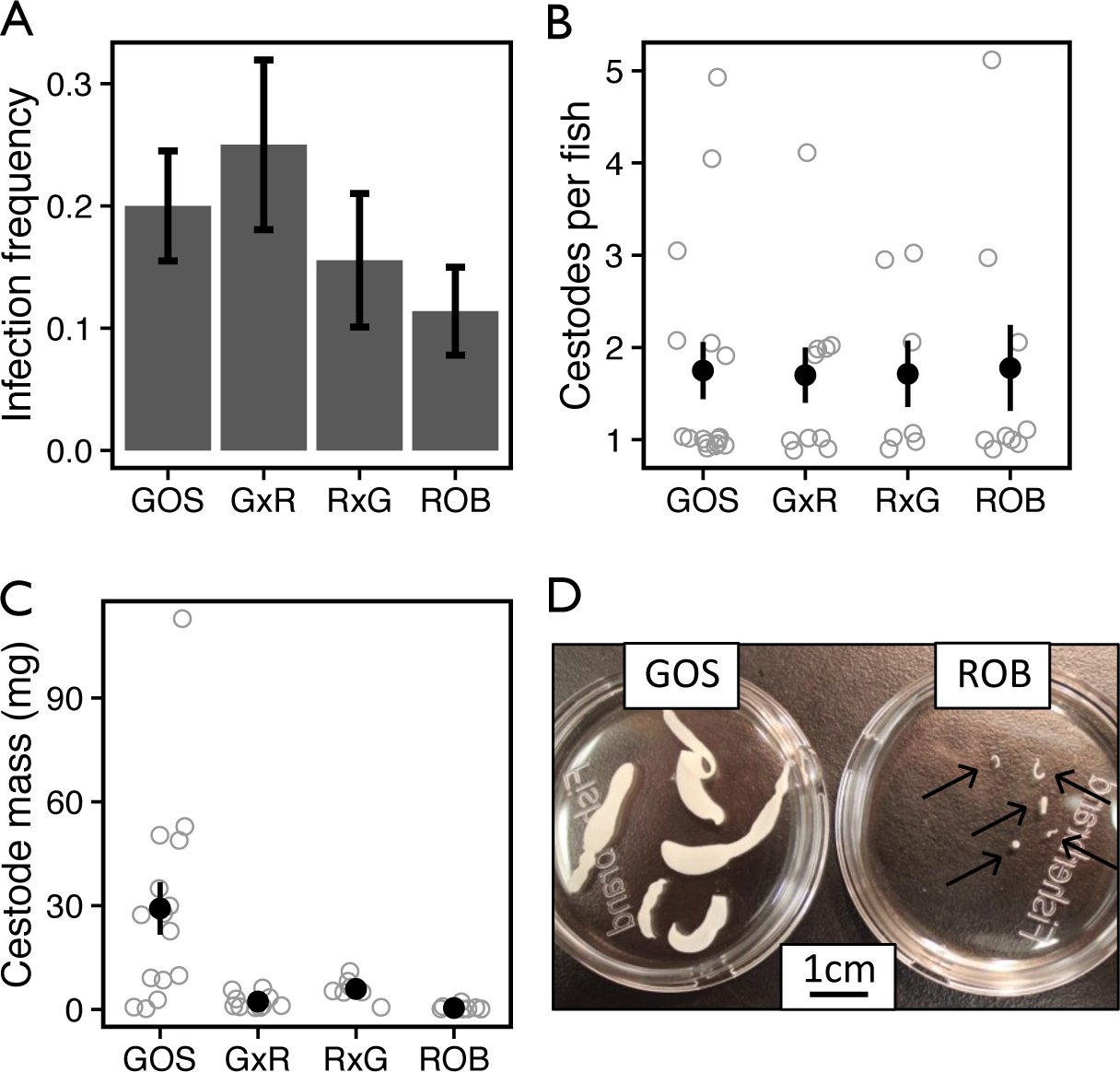
Infection success as measured by A) infection frequency (the proportion of stickleback with *S.solidus*), B) mean abundance of *S.solidus* per fish, and C) cestode mass (averaged across all cestodes in each fish). Error bars are SEs of population means. D) Full-sibling cestodes removed from GOS and ROB stickleback 42 days post-exposure. Fig. S7 presents the data in C) on a log-scale to better distinguish F1 hybrids from ROB, along with cestode masses from marine genotypes and F1 crosses between marine fish and GOS or ROB. Number of experimental fish per population in A): GOS=80), GxR=40, RxG=45, ROB=79. Number of infected fish per population noted in Fig 2.

Though infection rate and intensity were equivalent, full-sibling, age-matched cestodes grew 68-fold larger in GOS than in ROB stickleback. At the time of dissection, the mean mass of each *S. solidus* was 29.2mg (N=16, SE=7.3) in hosts descended from the high-infection population (GOS). In ROB fish, cestodes averaged only 0.4mg (N=9, SE=0.23). There was a significant host family effect on cestode growth, but no effect of cestode origin (Table S2). Cestodes grew to intermediate size in both GxR and RxG hybrids (means of 2.31mg and 5.9 mg respectively, with SEs=0.7 and 1.2; Fig3C). RxG hybrids had 2.5-fold larger cestodes than hybrids with GOS dams (t=2.589, P=0.019). Cestodes from GOS fish were 14-fold larger than those from GxR hybrids (post-hoc t-test P=0.064) and 5.6 fold larger than RxG hybrids (P=0.708). Conversely, cestodes from RxG hybrids were 11-fold larger than those from pure ROB fish (P<0.0112; GxR were 4.5-fold larger, P=0.114). Cestode mass was independent of host size, mass, condition (mass/total length), sex, or infection intensity. Note, however, that host mass would likely play an important role in older cestodes that are larger and more space-limited (22, 23).

Because most *S.solidus* growth occurs in stickleback, egg production is correlated with the adult parasite’s dry weight, and *S.solidus* must reach a minimum size (~10-50mg) to successfully infect its terminal host (24, 25), growth suppression effectively constrains the parasite’s reproductive potential. Arguably, this measure of parasite success represents an ‘extended phenotype’ (26) of the stickleback, in the sense that the host’s genotype alters the parasite’s phenotype. The intermediate size of parasites in hybrid fish also strongly suggests an additive genetic basis for this evolutionary difference.

To determine whether reduced parasite growth is an ancestral or derived trait, we measured *S.solidus* growth in susceptible marine fish (Weber et al. 2017). These cestodes grew quickly, reaching up to 320mg in 2.5 months (mean=130mg; Fig. S7). This is 4.5-fold larger than the mean size of *S. solidus* in GOS fish, possibly because the cestodes from marine fish were older (75-83 days for marine, 41-51 days for GOS). We then compared cestodes from pure marine fish with cestodes from marine x freshwater F1 hybrids (ROBxM and MxGOS, all assayed 2.5 months post-exposure). Consistent with the results reported above, cestodes from MxGOS fish (N=2) were 77.7-fold heavier than cestodes from the ROBxM fish (N=3; t-test, P=0.0011). Although this is not a large sample, the effect size is massive and quantitatively consistent with the fold-difference between cestodes from pure GOS and ROB fish. These data demonstrate that the growth-suppressive phenotype observed in ROB fish is an evolutionary gain of immune function from a susceptible and growth-permissive ancestral marine population, which has rapidly evolved over the ~12,000 years since Roberts Lake was likely colonized by marine fish.

### Growth suppression in Wild Populations

We hypothesize that the dramatic growth suppression observed in lab-raised ROB fish may explain the absence of observed infections in wild ROB fish. To test this hypothesis, we predicted that stickleback populations that more effectively suppress *S.solidus* growth would have correspondingly fewer observable infections. As expected, we found a positive correlation, across populations, between the mean size of *S.solidus,* and the parasite’s abundance. Focusing first on variation among fish within populations, we confirmed a well-established crowding effect (22, 23). Controlling for a random effect of lake, individual stickleback with more *S.solidus* have correspondingly smaller *S.solidus* (Fig.4A; mixed-model linear regression P<0.0001, controlling for host mass). However, we observed a counter-gradient trend across populations: populations with higher prevalence of *S.solidus* on average had larger cestodes (Fig. 4B, P=0.0058). GOS and ROB stickleback sit at opposite ends of this continuum, ROB growth suppression being so extreme that we could not locate infections in wild fish to measure cestode mass. This counter-gradient trend lends additional support to our inference that stickleback populations differ in their ability to suppress parasite growth, and that growth suppression is a repeatedly-evolved trait that reduces *S.solidus* prevalence. An important caveat, however, is that because these are natural rather than controlled infection, the age of each parasite is a potentially important but unknown covariate.

**Figure 4.**
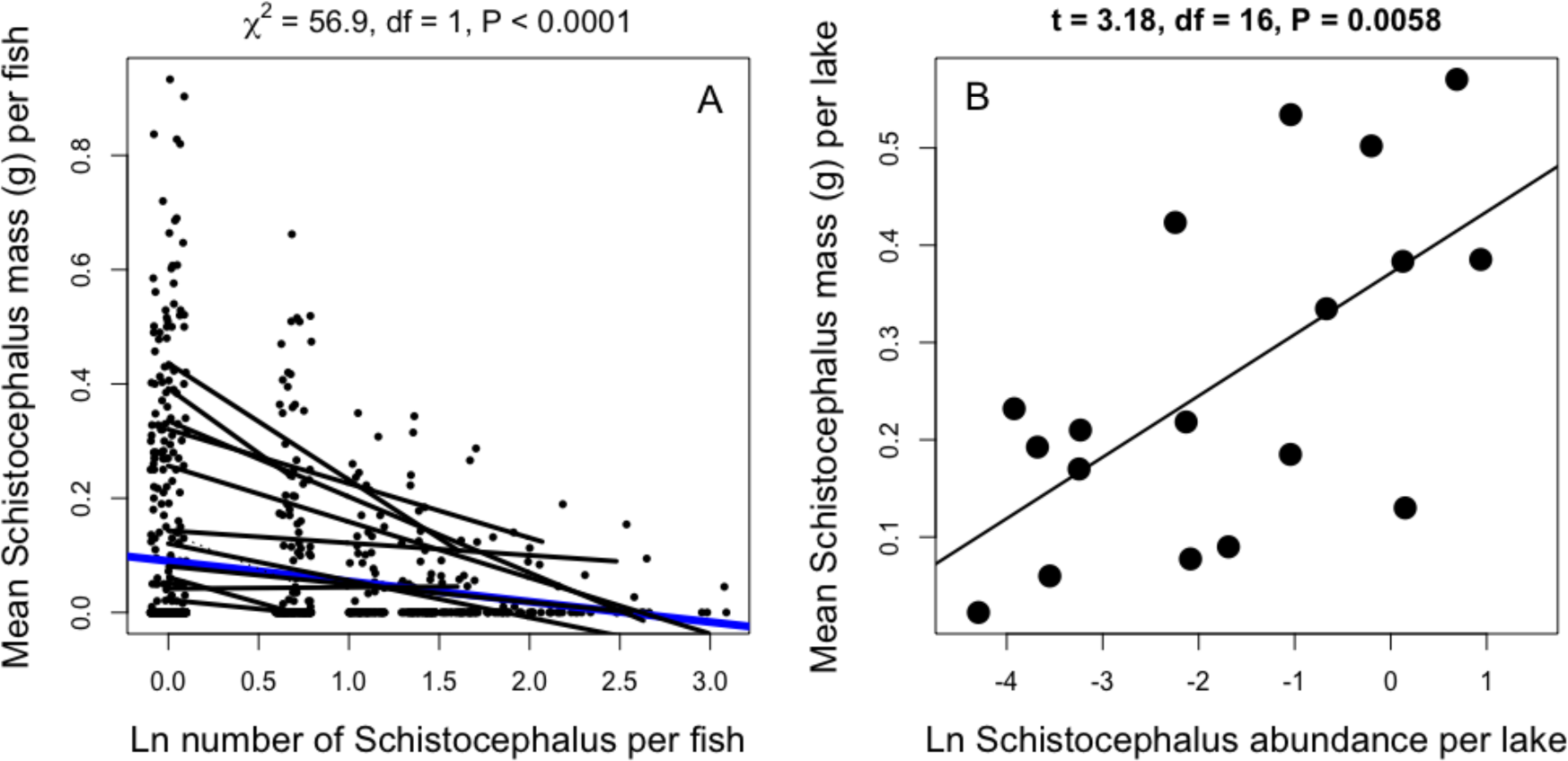
Cestode mass is correlated with infection abundance in the wild, but the effect direction is reversed when comparing A) individual fish within each of many populations, versus B) averages across populations. In A), points represent individuals within each of many populations. Within-population trendlines are shown in black, and consensus effect in blue (from a mixed model GLM). In B), we plot population mean cestode abundance versus mean cestode mass with lakes as the level of replication.

## Discussion

Stickleback are well known for their parallel evolution of adaptations to freshwater habitats (27, 28). This includes repeated independent evolution of increased resistance to freshwater-specialized parasites (18). However, as shown here, this immune evolution is not entirely parallel: ROB stickleback (but not GOS stickleback) evolved an ability to suppress *S.solidus* growth by two orders of magnitude. We infer that growth suppression by ROB fish explains the apparent absence of *S.solidus* in wild-caught fish from Roberts Lake. In laboratory trials, these fish are just as susceptible to infection as GOS fish. But, growth-suppressed cestodes can be so small that extant infections in wild-caught fish might be overlooked. Our survey of wild-caught fish suggests that growth suppression may be a widespread phenomenon; lakes with smaller *S.solidus* tended to also have lower parasite abundance.

To our knowledge this represents the first example of rapid evolution of a wild vertebrate’s ability to suppress helminth growth. However, several lines of evidence suggest that growth suppression may be common in other vertebrates as well. There is pervasive heritability for resistance-related traits in ruminant livestock (29). Most livestock studies measure helminth burden and resistance using non-invasive fecal egg counts (FECs). However, helminth length has very high heritability among hosts, and likely contributes to variation in FEC (30, 31). Even humans harbor heritable variation in helminth resistance and size (32). Notably, a study on hookworm weight in Papua New Guinea villagers suggests that humans vary in their ability to regulate parasite size.(33).

Bishop elegantly defined resistance as a host’s ability “to exert control over the parasite life cycle”, even in persistent infections [e.g., heritable variation for lower FEC (34, 35)]. Using this definition, stickleback control of cestode growth represents an effective form of resistance. In populations such as Gosling Lake that lack growth suppression, individual cestodes can grow to be up to 72% of the parasite-free mass of their host. Such large parasites can impose severe fitness costs by limiting a host’s ability to forage, grow, disperse, evade predators, and breed (19, 20, 36). To the extent that these effects depend on parasite size, growth suppression may partially or wholly rescue the fitness of ROB fish by facilitating survival or reproduction despite infection.

This benefit to the host can have ancillary effects on parasite epidemiology. In particular, smaller cestodes may be less fecund (25). Or, small cestodes may be less effective at manipulating host behavior in ways that may facilitate transfer to the terminal bird host (37, 38). Lastly, suppressing cestode growth may help the host clear the parasite infection. We occasionally observed small *S.solidus* encased in cysts (Fig. S8) within ROB but not GOS hosts. Some of these cysts contained degraded cestodes. We infer that growth suppression may aid in cyst formation (smaller parasites being easier to envelop), which in turn facilitates cestode-killing.

We also found heritable between-population differences in constitutive and infection-induced immune traits (granulocyte ROS production, and granulocyte frequency, respectively). Both traits have been linked to macroparasite killing by vertebrates (9, 39-42). But, we were unable to find reports of their influence on macroparasite growth, and our study is not designed to formally test this association. The infection-associated increase in HK granulocytes in ROB and RxG fish is consistent with the eosinophilia observed with helminthiasis in other vertebrate species (43). However, two observations suggest this is insufficient to explain variation in cestode growth. First, RxG hybrids had larger cestodes than ROB fish despite exhibiting comparable granulocyte abundance. Second, RxG hybrids had larger parasites than GxR hybrids, despite the former’s stronger granulocyte response. Although granulocyte ROS levels negatively correlated with cestode size (Figs. 2B&3C), this trend is confounded with overall population effects and does not hold among families within populations. The interactions between granulocyte-generated ROS and cestode growth warrant further investigation.

Although the proximate cause of cestode growth suppression remains unknown, our results offer several insights into stickleback immune evolution. First, even though GOS and ROB fish evolved to equally resist initial establishment by *S.solidus*, other immune traits exhibited non-parallel evolution. Relatively little is known about the extent of parallel (or, non-parallel) evolution of immune phenotypes across evolutionary replicate populations, in contrast to more easily-measured morphological traits. We suggest that parallel and non-parallel changes in immunity will prove to be useful in studying the context-dependency of immune evolution, and redundancy in immune function.

Second, we found that ROS production was unresponsive to *S.solidus* infection. This result is contrary to several previous studies, which are themselves conflicting. One study suggests that ROS response increases in early-stage *S.solidus* infections but is eventually suppressed (44), and another that granulocyte proportions fluctuate throughout infection but ROS levels only increase in late-stage infection (45). These different results may be a function of the particular populations studied (here Canadian, versus European stickleback and parasites used in prior studies) or variation in post-exposure time points of immune assays. Given the differences in granulocyte response between geographically nearby lakes (GOS versus ROB), it is quite plausible that host-parasite interactions on different continents may follow very different kinetics. Differences in ROS measurement methods may also play a role.

Finally, the maternally inherited increase in relative granulocyte abundance is an intriguing response that warrants additional attention. ROB but not GOS fish initiated a strong shift towards granulocytes following *S.solidus* infection, and F1 hybrids matched the response of their maternal parent. Many immunological maternal-effects in birds, fish, and mammals are related to antibody transmission between mother and offspring (46). These antibody-based effects tend to be relatively short-lived. Because fish in the present study were not exposed to *S.solidus* until adulthood, it is possible that a different mechanism may be involved.

Debate continues about the heritability of human immune responses (47, 48), but our results show that genetic variation for immunity segregates both within and among wild populations, and that immune differences influence parasite infection. Identifying the underlying mechanisms of natural infection variation will provide important contributions to this debate. Natural selection in disparate wild populations provides a powerful genetic scan to locate adaptive genetic variation involved in parasite resistance and immune function (49).

## Materials and Methods

### Estimating Natural Infection Prevalence

We used unbaited minnow traps to sample threespine stickleback (Scientific Fish Collection permit NA12-77018 and NA12-84188; Table S1). The majority of sites were sampled once in 2009 or 2013, but some were sampled repeatedly over multiple years (Table S1). All animals were euthanized by immersion in MS-222, fixed in 10% neutral buffered formalin, and then dissected to evaluate *S. solidus* presence.

### Breeding wild-caught stickleback

We generated crosses between wild-caught anadromous marine stickleback from Sayward Estuary to generate pure marine families (M genotype). We also generated one MxR cross and one GxM cross. Crosses were performed in June 2012 (Figure S1) and fertilized eggs were sent to the University of Texas at Austin (Transfer License NA12-76852), where all animals were reared in freshwater conditions and housed with full-siblings (9-39 animals per family) in either 40L or 10L tanks. All components of this experiment were approved by the University of Texas Institutional Animal Care and Use Committee (IACUC) as part of AUP-2010-00024.

### *S. solidus* Collection and Experimental Exposures

*S.solidus* were harvested from stickleback captured at three sites in British Columbia, Canada (Gosling Lake, Echo Lake, and Jericho Pond). The parasites were bred to obtain eggs from full-sibling families (22, 23), using media and methods described in (18, 50). Juvenile copepods *(Macrocyclops albidus)* were exposed to 1-2 freshly hatched tapeworm coracidia in 24-well plates. Two weeks after exposure we screened copepods to identify singly-and doubly-infected individuals, which were fed to fish between 21-26 days post exposure [when *S. solidus* coracidia are developmentally capable of infecting stickleback (51)]. Similarly aged, uninfected, copepods were used for control exposures.

Stickleback exposures began by isolating 5-7 fish per family and clipping either the first or second dorsal spine to distinguish control (1-2 fish per family) from experimental animals. At least 12 days after clipping, we moved individual fish into 1-liter containers and withheld food for 24 hours. Experimental and control fish were fed either 4-5 infected or uninfected copepods, respectively. The following day after confirming that all copepods were consumed, we combined animals into a single tank. 41-51 days post exposure (41-44 days for all but two families) we dissected fish, recorded fish metrics (length, mass, sex), noted the presence or absence of tapeworm infection, and performed immune assays. All fish were exposed to *S. solidus* between 15-23 months of age.

### Measurement of Immune Traits Via Flow Cytometry

We dissected head kidneys (HKs) from control and experimental fish and immediately placed organs in cold HK media (0.9x RPMI containing 10% FCS, 100 μM NEAA, 100 U/mL penicillin, 100μg/mL streptomycin, 55μM β -mercaptoethanol). HK were manually disrupted and filtered by grinding against a cell strainer (35μm mesh, Falcon 352235) and rinsing with 4mL of cold HK media. Cell suspensions were centrifuged at 300 g, 4°C for 10 minutes, the supernatant was removed, and the cell pellet was resuspended in the remaining media (~200μL). Live HK cells were counted using a hemacytometer (Hausser Scientific 3520) based on Trypan blue exclusion (Corning 25-900-CI). For each fish, HK cells were divided into three treatment groups: 1) media-only control, 2) DHR-123-stained, 3) DHR-123-stained and PMA-stimulated.

For each treatment group 2x10^5^ HK cells were plated in 200ul of HK media (group 1) or HK media containing DHR-123 (2 μg/mL; Sigma D1054) (groups 2 and 3). All samples were incubated in a 96-well plate at 18°C, 3% CO2 for 10 minutes. Next, PMA (130 ng/mL, Sigma P8139) was added to group 3 while the groups 1 and 2 received equivalent volumes of plain HK media. After incubating cells for an additional 20 minutes at 18°C, 3% CO2, all samples were run on an Accuri C6 Sampler and analyzed using FlowJo (Treestar). Granulocyte:lymphocyte ratios were calculated from the media-only controls (Fig. S2). The magnitude of ROS production was determined by comparing the median fluorescence of unstimulated and PMA-stimulated cells. Fig. S2 contains additional information on gating strategies.

### Statistical Analyses

All statistical tests were performed in the R software environment [V3.3-1; (52)]. For models without random effects, we performed Tukey HSD-corrected post-hoc comparisons using the multcomp package [V1.4-5; (53)]. We used the lme4 package [V1.1-12; (54)] to construct random effect models, and used maximum likelihood to calculate AIC scores for each model (Table S1). We used the lsmeans package [V2.23-5; (55)] to perform Tukey HSD-corrected posthoc comparisons on mixed models.

## ACKNOWLEDGEMENTS

We thank L. Ma for assistance with lab work. Wenfei Tong produced the illustrations in Fig. S1. Douglas Emlen provided comments on earlier drafts. This research was funded by the Howard Hughes Medical Institute.

---

**Figure S1.**
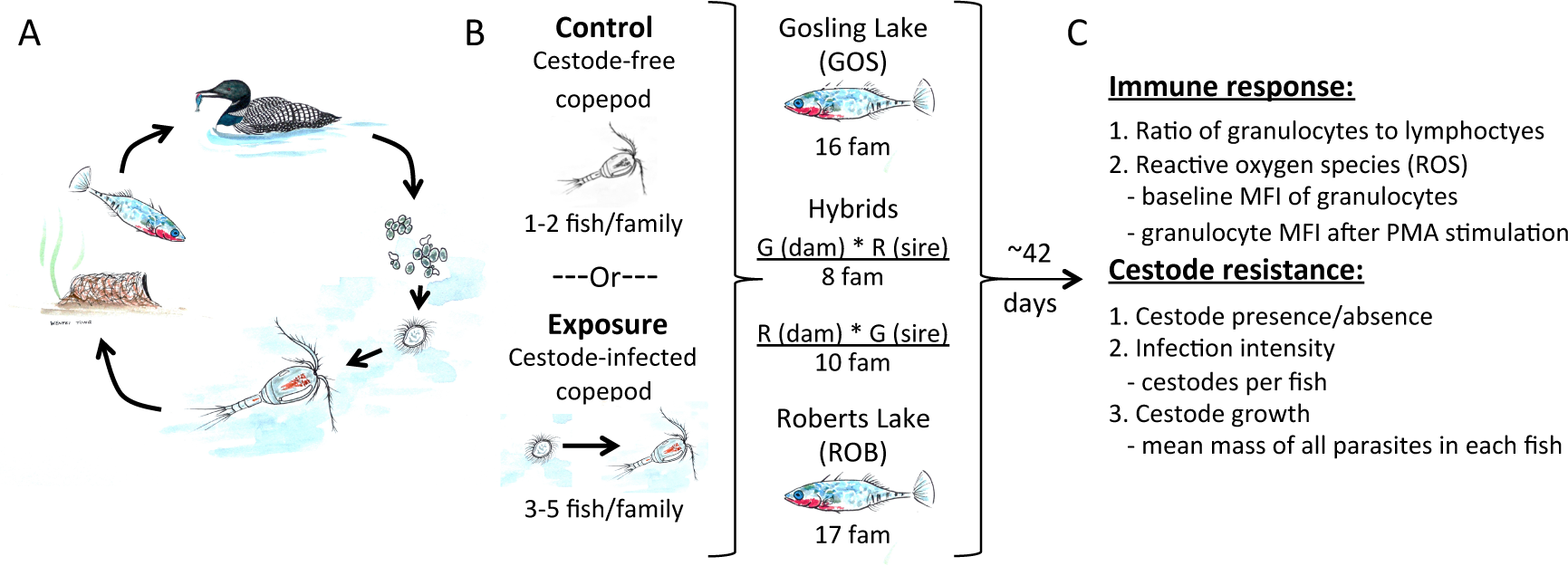
Experimental cestode infections. The life cycle of S. solidus (A) involves hatching from eggs in freshwater (i), a brief free-swimming period (ii), copepod infection (in), infection of threespine stickleback (iv), infection of picivorous birds and sexual reproduction (outcrossing or selfing) in this final host (v), followed by fecal excretion of eggs into water. (B) We fed (un)infected copepods to numerous lab-reared sticklebacks families (indicated as “fam”) to test whether fish from naturally high-and low-infection populations (and their hybrids) differ heritably in immune responses and parasite resistance. (C) A summary of immune traits and resistance traits measured on the control, exposed, and infected fish.

**Figure S2.**
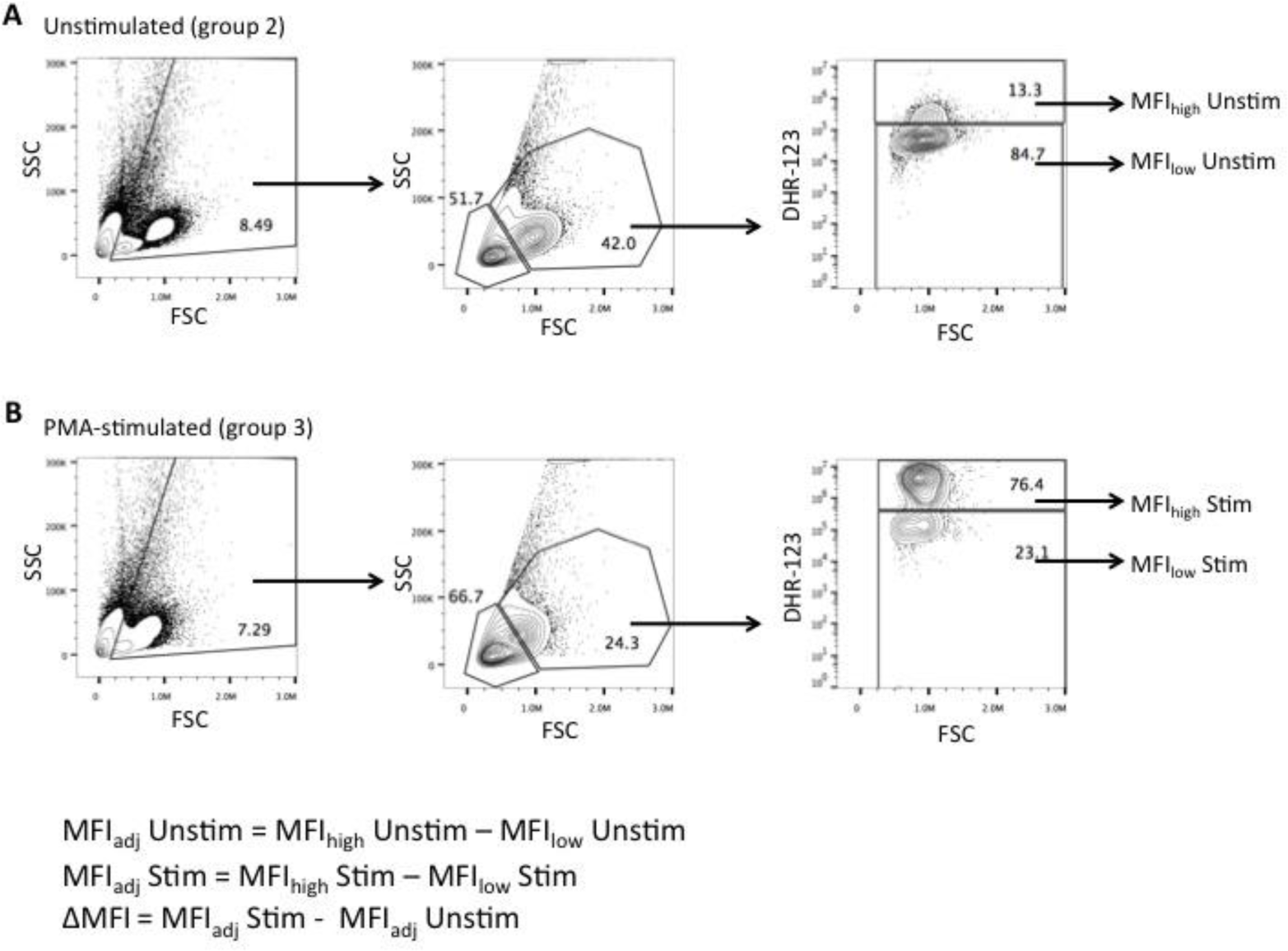
Calculation of median fluorescence intensity of DHR-123 stained HK granulocytes. The frequencies of HK leukocytes were determined via flow cytometry based on forward scatter (FSC) and side scatter (SSC) parameters. Granulocytes are FSC^high^ SSC^high^ while lymphocytes are FSC^low^ SSC^low^. The intensity of ROS production for each treatment group was determined based on median fluorescence intensity (MFI) of DHR-stained cells. To account for well-to-well and experiment-to-experiment variation in DHR staining intensity, we subtracted the the background DHR staining intensity (MFW population) from that of the DHR+ cells (MFIhigh) to obtain the adjusted MFI (MFI_adj_) for each sample. We calculated A) the background MFI of resting, unstimulated (MFI_adj_Unstim) and B) PMA-stimulated granulocytes (MFI_adj_Stim). The total change in ROS intensity (ΔMFI) was calculated by subtracting MFI_adj_Unstim from MFI_adj_Stim.

**Figure S3.**
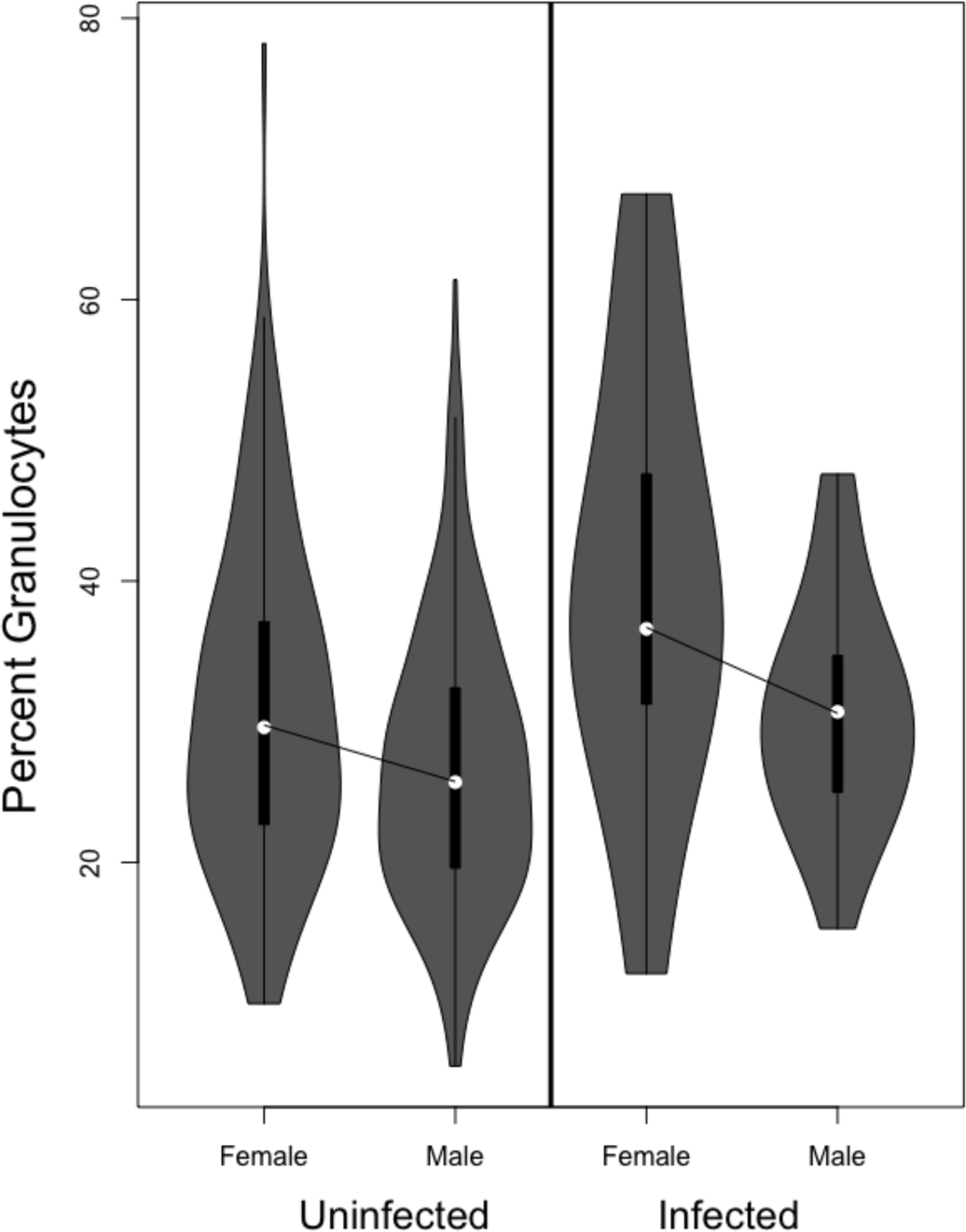
There is a significant but weak tendency for male stickleback to have relatively fewer granulocytes than females (or, equivalently, more lymphocytes). Because sex does not interact with fish or parasite genotype, here we lump together all host genotypes to focus on the sex effect in uninfected and infected fish, respectively.

**Figure S4.**
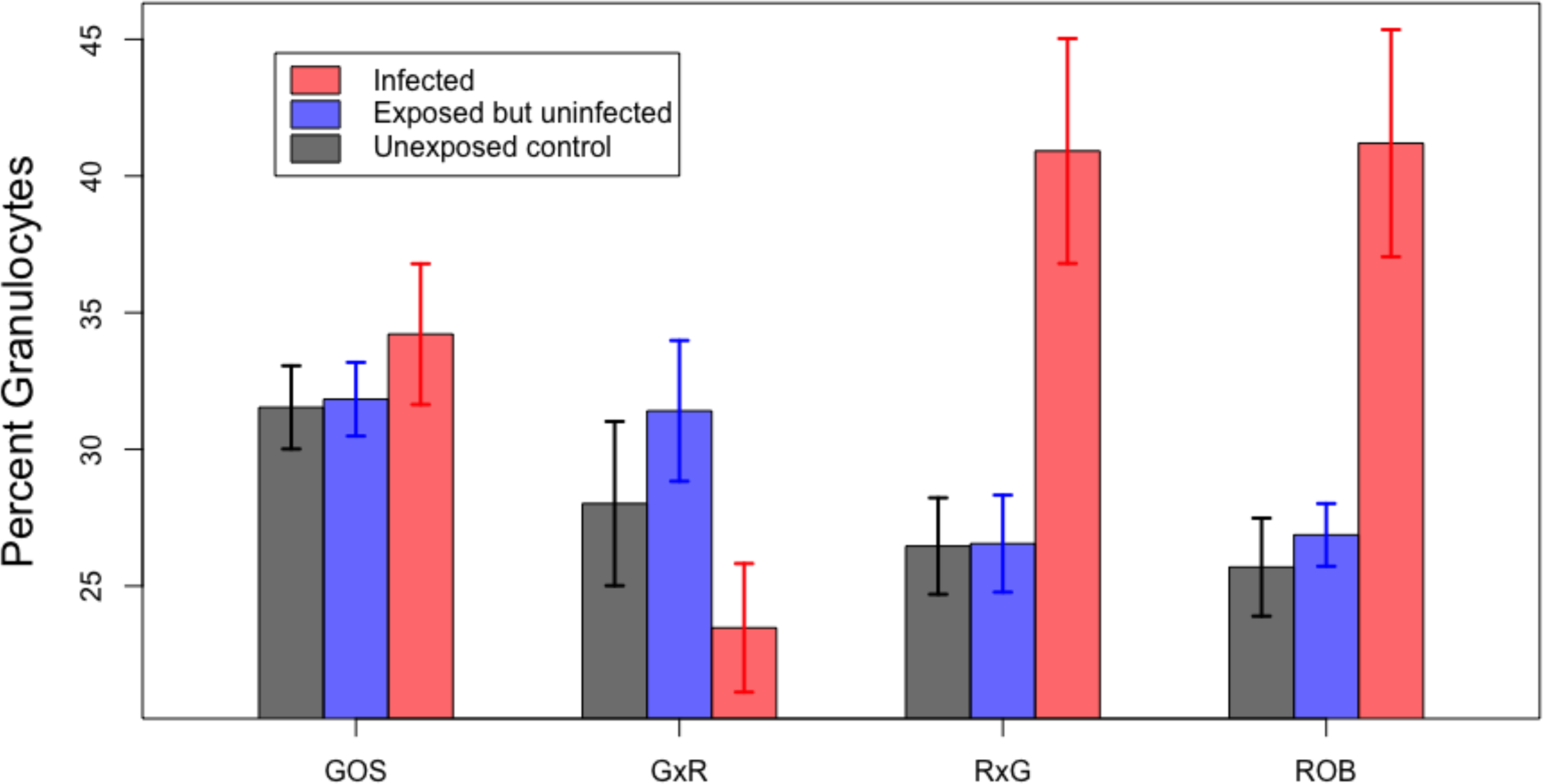
The percentage of granulocytes in HK cell culture, as a function of fish genotype (GOS, reciprocal F1s, ROB) and experimental infection status. Here, we distinguish between three infection states: unexposed control fish that did not encounter *S.solidus* (fed a copepod only), fish that were exposed but did not have a detectable infection 42 days later, and fish that were infected 42 days post-exposure. We observe no significant difference between unexposed and exposed-but-uninfected fish, so for the figures and results in the main text we combined these groups into a single ‘uninfected’ category.

**Figure S5.**
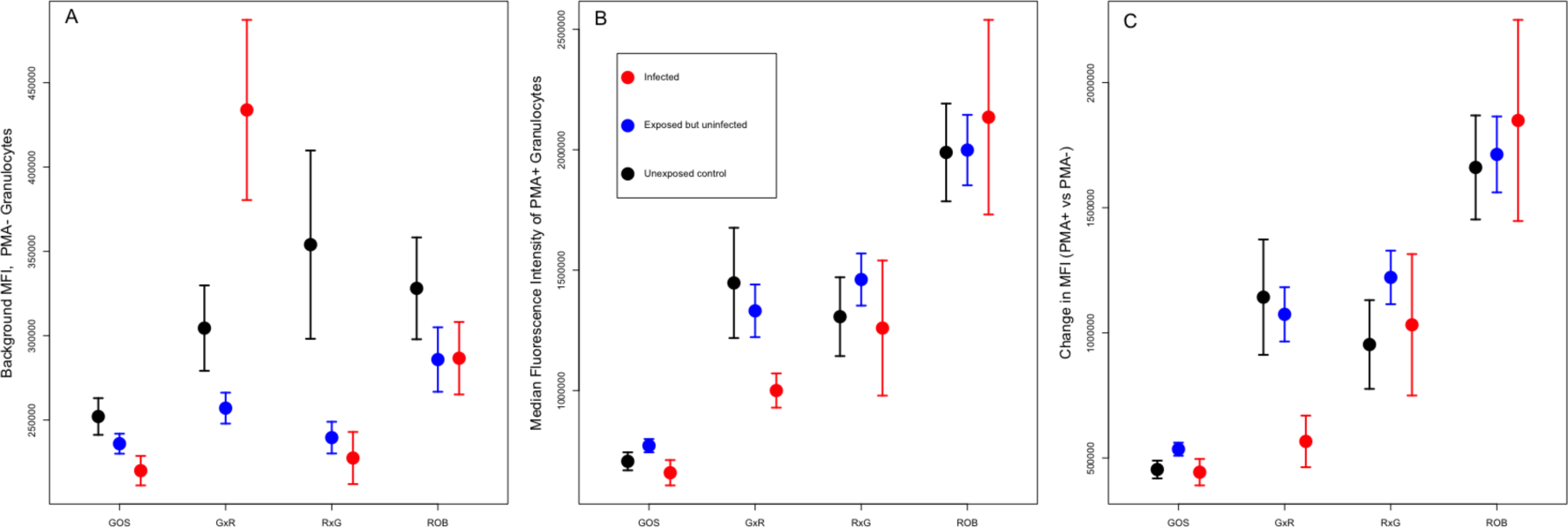
The effects of *S.solidus* exposure or infection, and host genotype, on ROS production as measured by the median fluorescence intensity (MFI) of granulocytes in HK primary cell culture. As in Fig. S3, we distinguish between unexposed control, exposed but uninfected, and infected fish. We measure three ROS variables here: (A) background MFI in resting state granulocytes (PMA^-^); (B) MFI following PMA stimulation, and (C) the difference between stimulated versus background MFI (ΔMFI). We observe no significant effect of *S.solidus* infection status, but a systematic trend for ROB genotypes to produce more ROS. Higher ROS in ROB fish is observed both for baseline ROS (A), and a stronger induced response to PMA stimulation of cell culture granulocytes (B,C). Reciprocal F1 hybrids are intermediate between the parental types, both for baseline and PMA-stimulated MFI, suggesting an additive genetic basis of this between-population difference. Baseline MFI varied significantly among families within populations.

**Figure S6.**
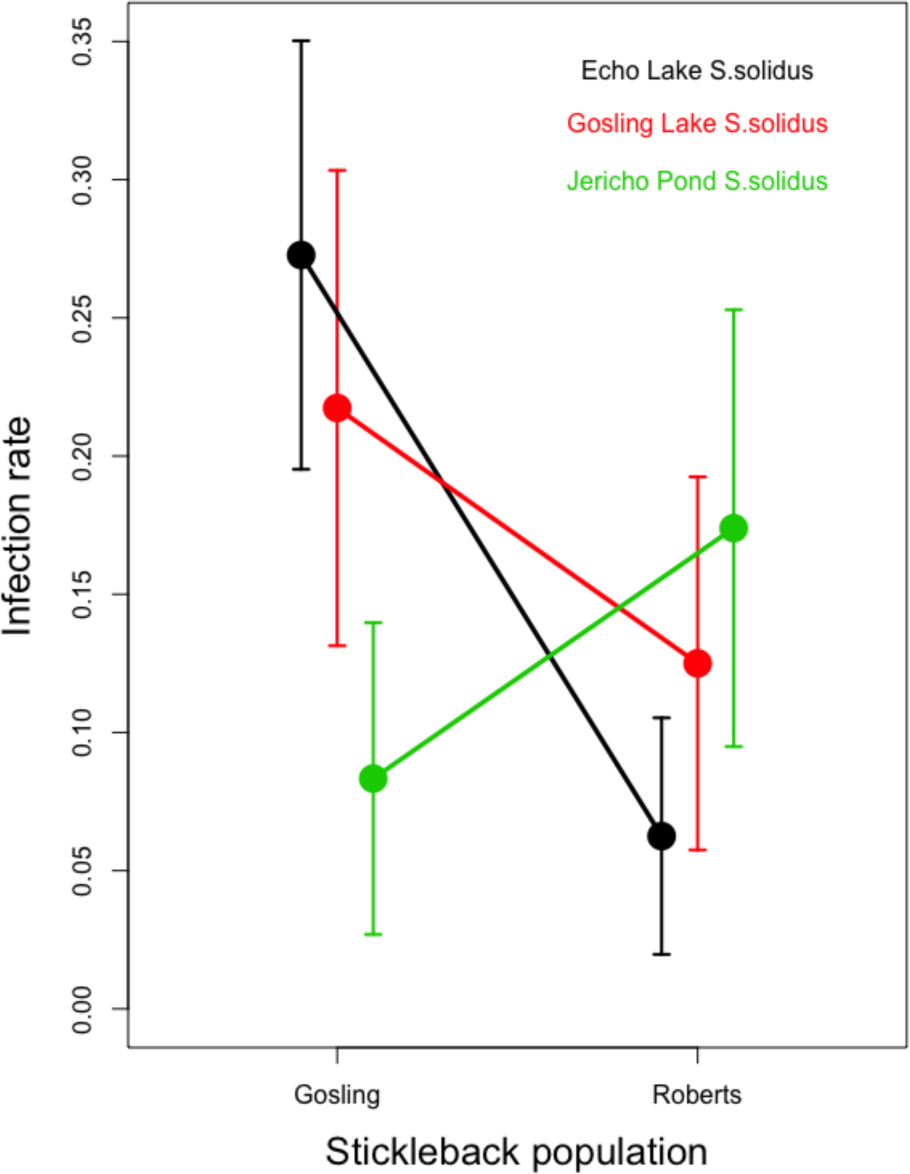
The effects of host and parasite genotypes on infection success rates. We observed a marginally non-significant interaction (P=0.09) between host and parasite genotypes, on the infection rates among experimentally exposed hosts. Here, we plot the infection rate as a function of host genotype (GOS or ROB)and cestode family. The geographically more distant cestodes, from Jericho Pond on the University of British Columbia campus, trended towards higher infection success in the otherwise-more resistant ROB fish, whereas both local cestode populations had somewhat higher success rate in GOS fish. Although no single contrast is significant, the different slopes for Jericho versus Echo or Gosling cestodes generates the marginally non-significant interaction.

**Figure S7.**
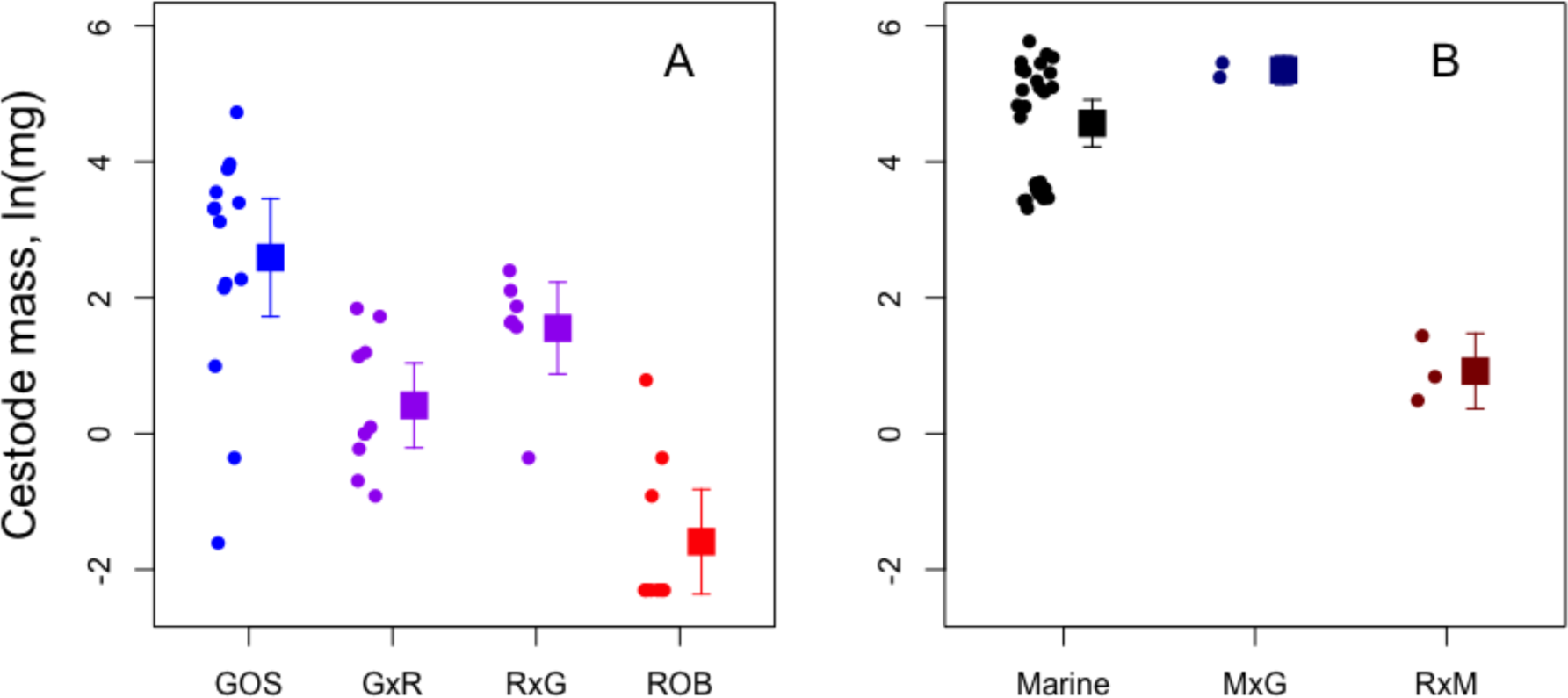
(A) Replotted data from Fig. 3C showing differences in cestode growth between lab-raised GOS versus ROB lake fish and their F1 hybrids, 42 days postinfection. Here we present cestode mass on a log-scale, to more clearly visualize the differences in masses of cestodes from F1 versus ROB fish. Each point is the mean cestode mass for an individual fish. The squares represent genotype average cestode mass with 95% confidence interval bars. GOS, ROB and F1 fish are plotted as blue red, and purple points,respectively. (B) Using the same log-mass y axis, we plot the mass of cestodes obtained from lab-raised pure Sayward Estuary marine stickleback (black points), and cestodes from F1 hybrids resulting from crosses of marine to Gosling Lake fish (dark blue) or marine to Roberts lake fish (dark red). In contrast to panel (A), these cestodes were measured after a longer infection period (75-83 days post-infection). Also, because we obtained relatively few infected fish, the data points in (B) represent individual cestodes, several of which come from an individual fish (e.g., the lower cluster of black points are 10 cestodes from a single host, and are correspondingly smaller than their less-crowded siblings). Post-hoc t-tests confirmed that cestode mass differed significantly between Marine versus MxR fish (P =0.00004), and between MxG versus MxR (P=0.0011).

**Figure S8.**
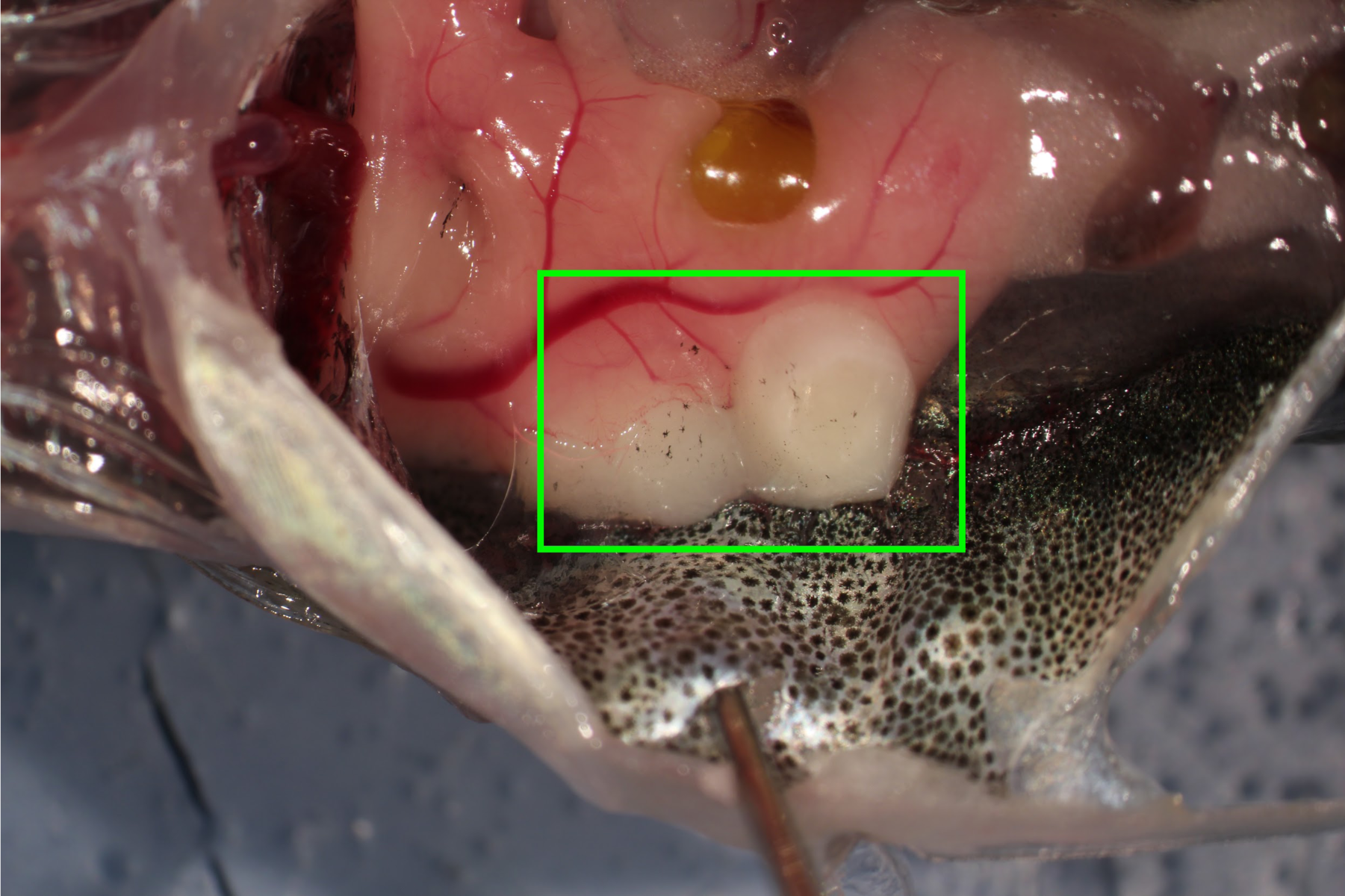
A *Schistocephalus solidus* cestode encased in a cyst (white structures within the green box), with the body cavity of a stickleback (head to left of image). Some cysts found in experimentally infected stickleback contained dead *S.solidus,* suggesting that small cestodes trapped in a cyst could be eliminated.

